# A simple and powerful analysis of lateral subdiffusion using single particle tracking

**DOI:** 10.1101/197152

**Authors:** Marianne Renner, Lili Wang, Sabine Levi, Laetitia Hennekinne, Antoine Triller

## Abstract

In biological membranes many factors such as cytoskeleton, lipid composition, crowding and molecular interactions deviate lateral diffusion from the expected random walks. These factors have different effects on diffusion but act simultaneously so the observed diffusion is a complex mixture of diffusive behaviors (directed, >Brownian, anomalous or confined). Therefore commonly used approaches to quantify diffusion based on averaging of the displacements, such as the mean square displacement, are not adapted to the analysis of this heterogeneity. We introduce a new parameter, the packing coefficient *Pc*, which gives an estimate of the degree of free movement that a molecule displays in a period of time independently of its global diffusivity. Applying this approach to two different situations (diffusion of a lipid probe and trapping of receptors at synapses), we show that *Pc* detected and localized temporary changes of diffusive behavior both in time and in space. More importantly, it allowed the detection of periods with very high confinement (~immobility), their frequency and duration, and thus it can be used to calculate the effective *k_on_* and *k_off_* of scaffolding interactions such those that immobilize receptors at synapses.

## Introduction

In cell membranes, molecules exhibit complex diffusive behaviours which reflect local heterogeneity of the membrane and/or interactions established with other molecules. Many factors such as the cytoskeleton, the lipid composition, crowding and molecular interactions affect diffusion simultaneously (1). The analysis of the transitions between different diffusive behaviours can provide hints about the organisation of the membrane and the interactions that a given molecule is undergoing. Single-particle tracking (SPT) fits well with this approach because it enables the localization of an individual molecule with nanometer precision, yielding detailed information on its motion (reviewed in (2)). However, the detection of transient changes in diffusive behavior along a trajectory is a long-standing problem for SPT techniques. Classical analyses based on the mean square displacement (*MSD*) and the calculation of the diffusion coefficient *D* are not appropriate due to the averaging intrinsic to their calculation (refs. in (3), (4)).

We introduce a new parameter, the packing coefficient *Pc*, which provides an estimate of the degree of free movement that a molecule displays in a period of time, independently of *MSD* and *D* calculations. *Pc* scales with the size of the confinement area; thus it is possible to identify periods of confinement by setting a threshold corresponding to a given confinement area size. Then it is possible to calculate the frequency and duration of confinement periods and to localize them in space.

An important type of interactions that membrane molecules may establish are scaffolding interactions, which are responsible for the accumulation of specific molecules in membrane domains. Scaffolding interactions immobilize for example receptors for neurotransmitters on the post-synaptic side of neuronal synapses (5). Due to the limited localization accuracy in SPT, immobilisation is translated into confinement in an area whose size is the localization accuracy. In this way, the Pc analysis is able to identify periods of transient immobilization. Assuming that these periods arise from a scaffolding interaction, the effective *k_on_* and *k_off_* of this interaction can be extracted from the frequency and the duration of immobilizations, respectively ((6) and references therein). This is particularly important as the understanding of molecular interactions under physiological conditions (*in cellulo*) is not straightforward. Classical bulk biochemistry used to identify interactions and to quantify molecular affinities favour the detection of strong molecular interactions. In addition, in these experiments molecules interact in an environment and under conditions that can be very different from the real situation in cells. This is particularly true in the case of reactions occurring in cell membranes. Therefore Pc analysis, by providing access to kinetic parameters of molecular interactions *in cellulo*, can help understanding the formation and dynamics of specialized membrane domains such as neuronal synapses within a living cell.

## Materials and Methods

### Cell culture and transfections

All animal procedures were carried out according to the European Community Council directive of 24 november 1986 (86/609/EEC), the guidelines of the French Ministry of Agriculture and the Direction Départementale des Services Vétérinaires de Paris (Ecole Normale Supérieure, Animalerie des Rongeurs, license B 75-05-20), and were approved by the Comité d’Ethique pour l’Expérimentation Animale Charles Darwin (licence Ce5/2012/018). All efforts were made to minimise animal suffering and to reduce the number of animals used. Primary cultures of rat hippocampal neurons were prepared as reported (7). For GFP-GPI experiments, neurons were transfected at 9 days in vitro (DIV) using Lipofectamine2000 (Invitrogen, Cergy Pontoise, France) following the manufacturer’s instructions. GFP-GPI plasmid was described elsewhere (7). SPT experiments were performed at 21-24 DIV. Neurons were transfected with SEP-γ2 ((8)) chimera at DIV 13-15 using Transfectin (BioRad, France) according to the manufacturer’s instructions. uPAINT experiments were performed one week after transfection.

### Drug treatment

Actin filaments were depolymerised with latrunculin A (3 μM; Sigma Aldrich, Lyon, France) solubilised in DMSO (Sigma Aldrich, Lyon, France). Cells were pre-incubated for 30 min with latrunculin or the control solution (0.002% DMSO), and used for SPT (performed in presence of the drugs). For the induction of cLTP, neurons were treated with glycine (200 μM) for 3 min in a bathing solution of osmolarity between 325 and 335 mosmol, containing (in mM): NaCl, 140; CaCl_2_, 1.3; KCl, 5.0; HEPES, 25; glucose, 33; TTX, 0.0005; strychnine, 0.001; and bicuculline methiodide, 0.02 (pH 7.4) (9) (all chemicals from Sigma Aldrich, Lyon, France). Cells were then rinsed and let in the bathing solution without glycine for SPT experiments. NMDA was applied for 3 min at 20 μM (10) (Tocris, UK).

### Single particle imaging with quantum dots

To track membrane protein diffusion, quantum dots (QDs) were pre-coupled to the corresponding primary antibodies, as reported previously (7). Briefly, goat anti-rabbit F(ab’)2-tagged QDs emitting at 655 nm (Q11422MP, Invitrogen, Cergy Pontoise, France) were incubated first with polyclonal rabbit anti-GFP antibody (132002, Synaptic Systems, Germany) or anti-GluA1 (AGC-004, Alomone Labs, Israel) for 30 min in PBS, and then blocked for 15 min with casein in a final volume of 10 μl. Neurons were incubated with the pre-coupled QDs (1:6000–1:10000 final QD dilution) for 5 min at 37°C.

All incubation steps and washes were performed at 37°C in MEM recording medium (MEMr: phenol red-free MEM, glucose 33 mM, HEPES 20 mM, glutamine 2 mM, Na-pyruvate 1 mM, and B27 1X). Cells were imaged within 30 min after QD staining. Neurons were imaged in MEMr at 37°C in an open chamber mounted on a IX70 inverted microscope (Olympus France, France) equipped with a 60X objective (NA 1.45; Olympus France). Fluorescence was detected using a xenon lamp, appropriate filters (QD: FF01-460/60-25, FF510-DiO1-25x36, FF01-655/15-25, Semrock, USA; GFP: HQ500/20, HQ535/30m; FM4-64: D535/50x, E590lpv2, Chroma Technology) and a CCD camera (Cascade 512BFT, Roper Scientific, France). QDs were recorded during 1000 consecutive frames (time points) at a frequency of 33 Hz (GFP-GPI) or 20 Hz (AMPAR).

### Single particle imaging with uPAINT

Neurons transfected with SEP-γ2 were imaged in MEMr at 37°C in an open chamber mounted on a N-STORM Nikon Eclipse Ti microscope equipped with a 100x oil-immersion objective (N.A. 1.49). Receptors were labelled with anti-GFP ATTO647N-coupled nanobodies (GFP-Booster, Chomotek). Nanobodies are small *Camelidae* antibody fragments (~15 kDa) consisting of a single monomeric variable antibody domain which allow specific labeling while introducing minimal linkage-error (11). After adding the solution of nanobodies to the medium (1:400 in PBS), cells were illuminated (622nm) with a laser (Coherent Genesis MXSLM, Coherent, France). Oblique illumination of the sample allowed to image nanobodies bound to their ligands in the cell surface without illuminating the molecules in the solution above ((12)). Fluorescence was detected using a C-NSTORM QUAD filter cube (Nikon) and an Andor iXon Ultra EMCCD camera (pixel size, 160 nm). SEP fluorescence was detected using a Nikon Intensilight lamp and appropriate filters (GFP Cube, Nikon; Ex 472/30 nm, DM495, Em 520/35). Nanobodies were recorded during 10000-20000 consecutive frames (time points) at a frequency of 33 Hz. The z position was maintained during acquisition using the Nikon perfect focus system. Mechanical x-y drift was corrected by adding individual fluorescent beads adsorbed on the glass coverslips, used as immobile references.

### Tracking and analysis of diffusion

Tracking was performed with homemade software (SPTrack_v4) in MatLab (The MathWorks, Natick, MA). The centre of the spot fluorescence was determined by 2D-Gaussian fit. Spatial resolution was ~10-20 nm for QDs and ~30 nm for nanobodies. The spots in a given frame (time point) were associated with the maximum likely trajectories estimated on previous frames of the image sequence. In uPAINT experiments, unbound fluorescent molecules freely diffusing in solution were discarded, their fast 3D diffusion avoided their detection in more than three images (trajectories of less than 10 points were discarded). Trajectories were defined as synaptic if they colocalized with a synaptic cluster. The mean square displacement (*MSD*) was calculated using

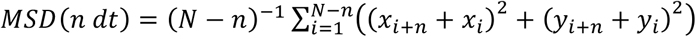

where *xi* and *yi* are the coordinates of an object on frame *i, N* is the total number of steps in the trajectory, *dt* is the time interval between two successive frames, *n* is the number of frames and *ndt* is the time interval over which displacement is averaged. The diffusion coefficient *D* was calculated by fitting the first 2 to 5 points of the *MSD* plot versus time (refs. in (13)) with the equation

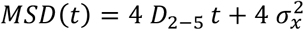

with σ_*x*_ the spot localization accuracy (positional accuracy, (2)) in one direction.

### Packing coefficient

The packing coefficient (Pc) at each time point *i* was calculated as

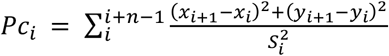

where *x_i_, y_i_* are the coordinates at time *i*; *x_i+1_*, *y_i+1_* are the coordinates at time *i+1, n* is the length of the time window (when using QDs, *n*=30 time points; in uPAINT experiments, *n*=10 time points) and *S_i_* is the surface area of the convex hull of the trajectory segment between time points *i* and *i+n*. *S_i_* was calculated using the *convhull* function in MatLab (MathWorks) (Fig. 2A). The validity of the packing coefficient to detect transient stabilization periods was checked on Monte Carlo simulations of brownian or confined trajectories.

### Monte Carlo simulations

The program was written in MatLab (The MathWorks, Natick, USA) and run on a personal computer (Dell Precision T1700). Trajectories were simulated as in Renner et al. (14), with some modifications. The *x* and *y* components of the *i-th* displacement step in the trajectory were randomly selected from two independent normal distributions with the mean of zero and the variance equal to 2*D_sim_ Δt*, using different *Δt* and *D_sim_* as indicated (Fig. 2C). The noise introduced in SPT trajectories by the limited accuracy of localization was simulated by adding a distance to *x* and *y*, distance chosen from an independent normal distribution with the mean of zero and a given variance that corresponds to the desired localization accuracy. This distance was calculated independently for *x* and *y* at each time point. Typically, trajectories had one or more periods of confinement, in which the positions were forced to stay in within a circle of the selected diameter *L* and reflecting border. Periods of confinement were imposed by inserting a boundary at a given time during the run, and removing it at a later prescribed time. To analyse the effect of acquisition frequency, trajectories were simulated with a *Δt* that matched the chosen acquisition frequencies.

### Statistical analyses

Statistical analyses were done using two-tailed Student’s t test, Mann-Whitney test (MW), one-way ANOVA or Kruskal-Wallis with Dunn multiple comparison test (KW) using Prism (GraphPad software, USA). Images were prepared using Photoshop (Adobe Systems, USA).

## Results and discussion

### The averaging effect of MSD calculation overlooks transient confinement periods

The *MSD* provides the simplest type of classification of the diffusion behaviour. Brownian motion yields a *MSD* = 4*D*τ (in two dimensions) where *D* is the diffusion coefficient and τ the time interval. In contrast, anomalous diffusion is characterized by a non-linear *MSD*=4*Dτ^α^* with *α* < 1. When diffusion is confined, the *MSD* asymptotically approaches a value related to the size of the confinement area (15,16). Unfortunately, the information obtained from *MSD* analysis is limited and, in practice, different situations of non-Brownian diffusion can produce the same *MSD*. This is due to the fact that the *MSD* is calculated by averaging all the displacements within the trajectory that correspond to a given time interval (see Materials and Methods section). Thus, if the molecule switches between different diffusive behaviours, the final *MSD* depends not only from the difference in diffusivity but also from the duration of each behavioral (brownian/anomalous/confined) period.

Fig. 1 illustrates some of the caveats of *MSD* analysis. We simulated random walk trajectories (10s-long with *Δt* =10 ms, see Materials and Methods) without (A1) or with a period of confinement that lasted 3 s in an area of 50 nm in diameter (A2) or that lasted 5 s in an area of 100 nm in diameter (A3). A fourth trajectory (A4) was always confined in an area of 100 nm in diameter. The *MSD* (B) of the trajectory A4 shows its confinement. In case of trajectories A2 and A3, the MSD plot (B) displayed the expected shape of confined diffusion, however, it should be noted that both trajectories had the same *MSD* despite their different confinement (3s in a compartment of 50 nm in diameter for A2, 5s in a compartment of 100 nm in diameter for A3, Fig. 1B). Next, we simulated trajectories that switched between periods of free diffusion (*D_sim_*=0.02 μm^2^/s during 3.75 s) and strong confinement (corralled in a 30 nm-diameter area during 22.5s) (Fig.1C). We chose three examples with confinement periods that were distanced (C1), apposed (C2) or co-localized (C3). The calculation of *MSD* on these three trajectories (Fig. 1D) allowed to detect the confinement of trajectory C3 and the “hopping” behaviour of trajectory C2. However, the trajectory in C1 produced a *MSD* similar to that of a trajectory that displayed slower free random movement (A1, Fig. 1). Thus, the *MSD* hid the confinement of trajectory C1.

**Fig. 1:**
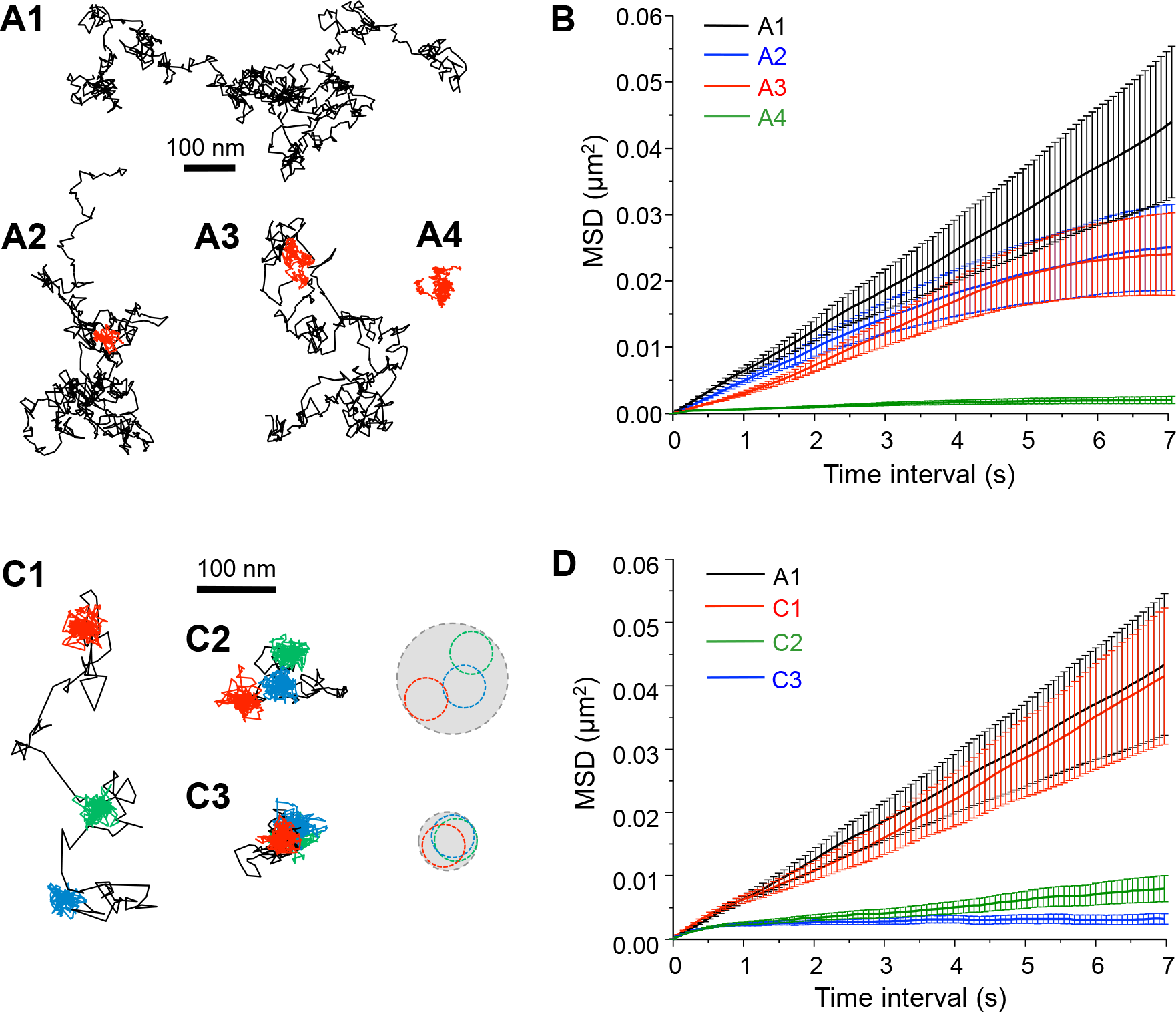
The averaging effect of *MSD* calculation within the trajectory overlooks transient confinement periods. (A) Brownian simulated trajectories without (A1), with one period of confinement (in red) of 3 s in a confinement area of 50 nm in diameter (A2), with one period of confinement of 5 s in a confinement area of 100 nm in diameter (A3) or always confined in an area of 100 nm in diameter (A4). (B) *MSD* values of the simulated trajectories in (A) (MSD ± s.e.m.). (C) Brownian simulated trajectories with three periods of confinement (in color) in 30 nm diameter areas that were distanced (C1), apposed (C2) or co-localized (C3). On the right: minimum circles containing each confinement period (in colors) or the whole trajectory (grey). (D) *MSD* values of the simulated trajectories in (C) and of trajectory A1 for comparison (MSD ± s.e.m.).

In order to identify and quantify transient changes between free and confined diffusion we set a new diffusion parameter, the “packing coefficient” (*Pc*). *Pc* quantifies the degree of “compaction” of a trajectory by comparing the length of the trajectory in a short time window and the surface area that it occupies (both to the square):

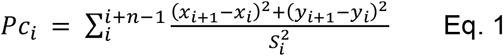

where *x_i_, y_i_* are the coordinates at time *i*; *x_i+1_*, *y_i+1_* are the coordinates at time *i+1, n* is the length of the time window and *S_i_* is the surface area of the convex hull of the trajectory segment between time points *i* and *i+n*. (Fig. 2A, see Materials and Methods). Unless indicated, we set *n*=30 time points (see below).

**Fig. 2:**
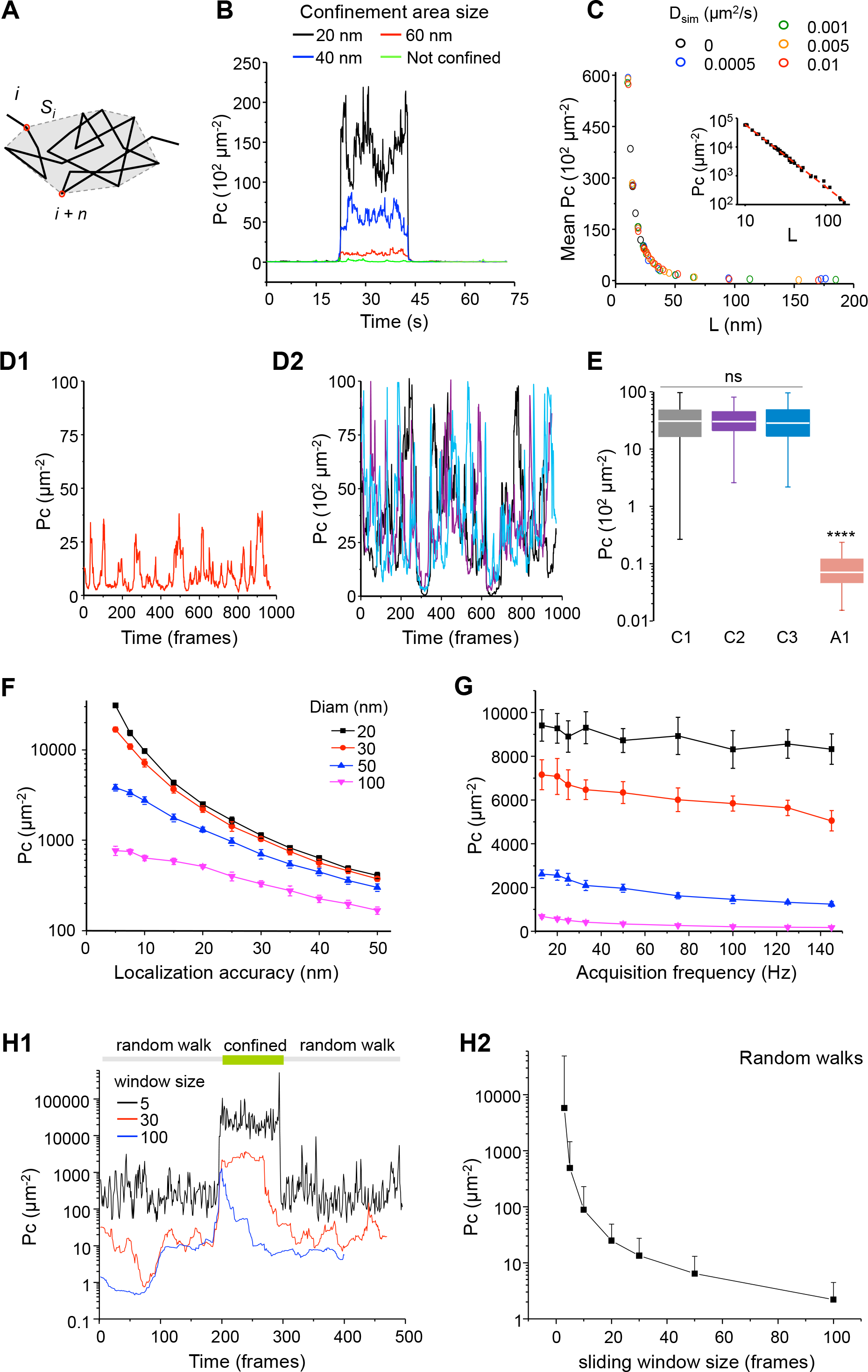
Detection of transient confinement using the packing coefficient (*Pc*). (A) Scheme representing the calculation of *Pc* at time point *i. S_i_* is the convex hull of the trajectory segment *i-i+n*. The sum of the square displacements between successive time points of the trajectory stretch spanning from *i* to *i+n* is divided by *S_i_^2^*. (B) *Pc* values of examples of 75 s-long simulated trajectories that were fully Brownian (green) or that have a period of confinement in areas of the indicated sizes. (C) Mean *Pc* vs the diameter of the confinement area *L* on simulated trajectories constructed with the indicated D_sim_. Insert: the same plot in log-log axis (D) *Pc* values of the simulated trajectories A1 (D1, always Brownian) and C1-3 (D2, with immobilization periods; C1 in black, C2 in purple, C3 in blue) of Fig. 1. (E) *Pc* values for the trajectories in (D) (median, 25%-75% IQR), Kruskal-Wallis test with Dunn’s multiple comparisons ns: not significant, ****: p<0.0001). F) Mean values of *Pc* during confinement periods of the indicated sizes, obtained on trajectories simulated with different localization accuracies (mean ± s.e.m, n= 1000; note the semi-log scale). G) Values of *Pc* during confinement periods of the indicated sizes, obtained on trajectories simulated with a localization accuracy of 10 nm and an interval of time between trajectory points that matched the desired acquisition frequencies (see materials and methods). (H) Effect of the size of the sliding window. (H1) *Pc* values of simulated trajectory with a confinement period of 100 time points. *Pc* was calculated using different sliding window sizes. The largest windows (100 time points) could not detect properly the confinement period. The shortest window (5 time points) accurately detected this period but the fluctuations were significantly higher than for longer sliding windows (note the semi-log scale). (H2) *Pc* values (mean ± s.e.m, n= 1000) obtained on Brownian simulated trajectories using different sliding window sizes. The statistical uncertainty increases in shorter windows.

In order to evaluate the capacity of *Pc* to detect transient confinement periods, we first simulated trajectories undergoing simple Brownian diffusion with or without a transient confined period in areas of variable sizes (Fig. 2B-C). *Pc* values were higher when the confinement area was smaller (Fig. 2B). Importantly, when simulations were done also varying the global *D* (*D_sim_*) of the trajectories, *Pc* scaled inversely to the size of the confinement area *L* independently of *D_sim_* (Fig. 2C).

Theoretically *Pc* tends to infinite when *L* tends to zero. However in case of trajectories obtained experimentally, the localization accuracy imposes a limit to the minimum *L* that can be observed. Thus we simulated trajectories taking into account the localization accuracy (see materials and methods). Under this conditions, the relationship between *Pc* and *L* followed the expected power law (inset Fig. 2C) with

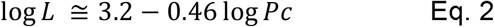

As shown in Fig. D1 and D2, *Pc* versus time for trajectories of Fig. 1 C (that underwent analogous confinement periods) were similar for the three trajectories (C1-3) but different from that of the trajectory A1 (free diffusion) (Fig. 2E). Thus *Pc* correctly detected and quantified the temporary changes in diffusion behaviour in these trajectories.

As expected, localization accuracy affected the capacity of *Pc* to evaluate the confinement. The relative differences in *Pc* due to different sizes of confinement were smaller when the localization accuracy decreased (Fig. 2F). *Pc* values also varied with the acquisition frequency but this did not preclude the detection of confinement. Confinements in circles of 20-100 nm in diameter were distinguishable with sampling frequencies between 13 and 145 Hz (Fig. 2G) and the relationship between *Pc* and *L* (eq. 2) was conserved. Finally, the length of the sliding window had to be short enough to optimize the ability to detect short stabilization events (Fig. 2H1) but long enough to sense the confinement and to decrease statistical uncertainty (Fig.2H2). Given our acquisition frequency, we set this value to 30 time points.

Many methods have been described to characterize various diffusion behaviours, each one fitting well with a particular experimental situation (for a review of analytical approaches, see (17)). The *Pc* analysis presents some advantages with respect to other strategies particularly in case of populations of molecules diffusing slowly and displaying multiple periods of confinement. The method proposed by Simson et al. (18), developed initially to study the effect of lipid rafts on lateral diffusion, is commonly used to detect confinement by calculating a confinement probability level comparing local and global diffusivity. This method allows identifies periods in which the molecule remains in a region for longer duration than predicted by a Brownian motion model. This is done by using a sliding time window to calculate the local maximum displacement and comparing it to the global *D* (*D* calculated over the entire trajectory). However, this method has two drawbacks: 1) random walks can be discriminated from confinement periods only if the presence of confinement does not affect global *D* and if the difference between the diffusivity of the random and confined trajectories is large enough (Fig. S1 in the Supporting Material) and 2) to access to short-lived changes in diffusion the calculation of *D* has to be performed on short trajectories segments that render the calculation of *D* less reliable (3,4,19). An alternative and derived approach compares *D* to the variance of displacement steps (20). However this strategy can only be successfully applied, once more, if the difference in the diffusivity between random and confinement states is large. The gyration quantification method (21) determines the area a given molecule explores by computing the radius of gyration. It is an efficient approach, but only if the trajectories have enough steps to define the motion (~50 steps). Furthermore, the fast and slow diffusion coefficients should differ by at least a factor of 5 (21). In contrast, in our experiments we found that *Pc* is independent from the global diffusivity and it efficiently discriminated confinement from slow Brownian movement (Fig. S1 in the Supporting Material).

Other proposed approaches analyse, e.g., the autocorrelation function of squared displacements (22) or the first passage time variance (23). Yet, in these two cases the temporal and spatial positions of the confinement periods are not provided. In contrast, the analyse of *Pc* vs time allowed the localization of the confinement periods that could then be correlated to a specific membrane region such as neuronal synapses (see below).

The *Pc* analysis as well as others confinement analysis face a main problem: Brownian diffusion trajectories can temporarily mimic confinement due to random fluctuations of the length of the displacements. However, the amplitudes and durations of these fluctuations are most of the time smaller and shorter than the ones associated with real non-Brownian transient motion (18). Therefore, the use of a threshold value of *Pc* (*Pc_thresh_*) and of a minimal duration above this threshold (*t_thresh_*) can suppress the detection of apparent non-random behaviors without excluding the detection of real confinement. *P_thresh_* can be set to the P95 or P99 percentile of *Pc* distribution of simulated random walks that are matched to the experimental data in acquisition frequency, localization accuracy and length of trajectories. The duration time threshold *t_thresh_* can then be chosen by applying *P_thresh_* to random walk trajectories and extracting the P95 or P99 of the distribution of durations. Thus, true confinement corresponds to a period with *Pc* being above *P_thresh_* that lasts more than *t_thresh_*. For example, when *Pc* was calculated over a window of 30 time points on random walks, P95 was equal to 67 μm ^−2^ (Fig. S2). Using this value as *Pc_thresh_*, we found *t_thresh_*=0.81s (P99 of the distribution of durations). Alternatively, the definition of a threshold that corresponds to a given confinement area size *L_thresh_* allows identifying the time and position in space of the trajectory sequences confined in areas of size ≤ *L_thresh_*. In this case, *t_thresh_* will depend on the acquisition frequency and the characteristic time of confinement.

### Detection of diffusive behaviour transitions on GFP-GPI trajectories

Membrane proteins and some lipids may undergo “hop diffusion”, being temporarily confined in 30–700 nm-diameter compartments and displaying frequent jumps between adjacent compartments (24). In cultured CHO cells, GPI-anchored GFP (GFPGPI) molecules display hop diffusion between compartments of about 40 nm in diameter, the size of which depend on F-actin integrity (24, 25).

We first tested the *Pc* analysis on SPT trajectories analyzing the diffusion of GFP-GPI on cultured hippocampal neurons. GFP-GPI molecules were labeled with quantum dots (QDs) coupled to an anti-GFP antibody (QD-GFPGPI). QD-GFPGPI were detected with a localization accuracy of 20-30 nm and tracked at 33 Hz. QDs blink, thus in order to avoid introducing unwanted uncertainty *Pc* was calculated only if the dark state within the sliding window was less than 5 time points-long.

As exemplified in Fig. 3A1-2, *Pc* values versus time for each trajectory revealed various patterns. Most trajectories had low *Pc* values characterizing free diffusion (Fig. 3A1) whereas some displayed high *Pc* values suggesting constrained (not free) diffusion (Fig. 3A2).

**Fig. 3:**
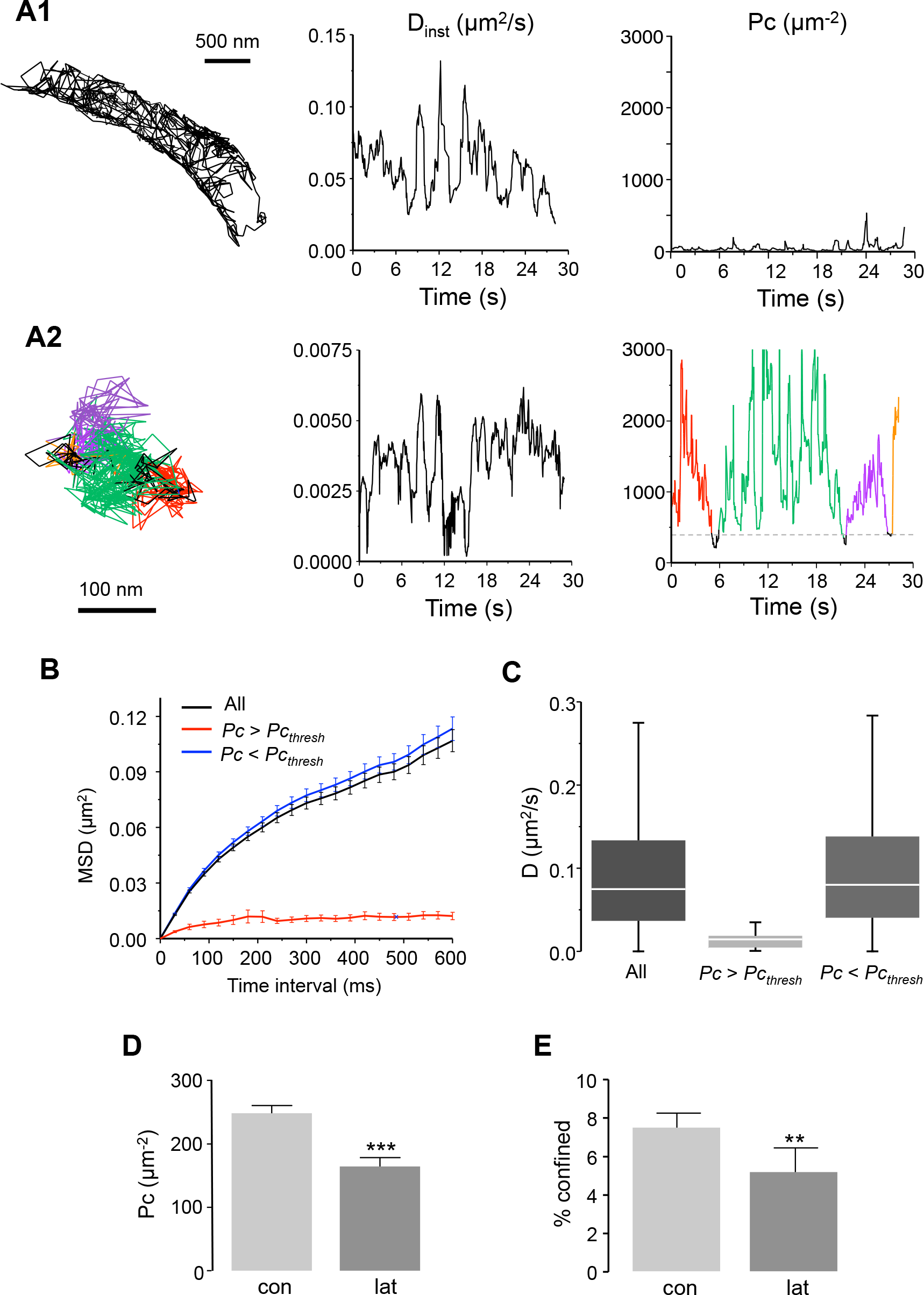
Lateral diffusion of GFP GPI on neurites, effect of F-actin depolymerisation. (A) Examples of QD-GFPGPI trajectories (left), with the corresponding instantaneous *D* (D_inst_, center) and *Pc* (right) plots. D_inst_ and Pc were calculated on the same sliding window (30 time points). (A1) trajectory without confinement. (A2) trajectory with multiple periods of confinement in areas ≤100nm (*Pc_thresh_*=406 μm^−2^). The detected periods are shown in color. (B) Average *MSD* plot of all QD-GFPGPI trajectories (black), portions of trajectories with *Pc* below (*Pc*≤*Pc_thresh_*, blue) or above (*Pc*>*Pc_thresh_*, red) threshold (n=275-926 trajectories). (C) Distribution of *D* (median, box: 25%-75% IQR, whiskers: 5%-95%) for the trajectories in (B). (D) Mean *Pc* values of trajectories in control condition (con) or alfter latrunculin treatment (lat) (mean+/- s.e.m., Mann-Whitney test ***: p<0.001, n=40-70). (E) Percentage of confined trajectories (C1), in control conditions (con) and after latrunculin application (lat) (mean+/-s.e.m., t-test **: p<0.01, n=40-70).

In order to quantify the diffusive behaviours of QD-GFPGPI, we set an initial *Pc_thresh_* of 406 μm^−2^, which corresponds to a confinement area size of 100 nm. In Fig. 3A2, the portions of trajectories for which *Pc* values are above *Pc_thresh_* (thus confined) appear in different colors. The mean *Pc* value during these periods was 2120 ± 125 μm^−2^, which corresponds to a confinement area size of 44.87 ± 3.31 nm in diameter. Interestingly, this value is close to the proposed size of hopping confinement areas described by Umemura et al. for GFPGPI (40 nm, (24)). However in our experiments, using a much lower acquisition frequency (33Hz) than the one used by Umemura et al. (50 kHz), only 7.50 ± 0.75 % of the trajectories displayed this behavior (24). Thus, in our recordings we detected the corralled molecules that remained a sufficiently long time in a hopping compartment to be detected at 33 Hz of sampling frequency.

We then sorted portions of QD-GFPGPI trajectories in two groups: those with *Pc* > 406 μm^−2^ (“confined”) and those with *Pc* <= 406 μm^−2^ (“free”). In each case, we analysed their diffusion comparing their *MSD* and *D* (Fig. 3B-C). Further, we compared the results obtained with the intact trajectories (“whole”) with those obtained on “free” and “confined” sections of trajectories. The *MSD* and *D* of the “whole” trajectories were close to that of the “free” group, overlooking the presence “confined” events (Fig. 3 B-C).

The disruption of the actin cytoskeleton using latrunculin increases the mobility of GFPGPI (24). This effect was attributed to a reduction in the amount of F-actin bound “pickets” that act as obstacles to diffusion and ultimately define the hopping compartments (26). We then checked if the confinement highlighted by Pc could be modified by F-actin depolymerization. As expected, the *Pc* values were significantly lower under the latrunculin condition (Fig. 3D) and the percentage of confined trajectories decreased by 30% (control: 7.50 ± 0.75 %, latrunculin: 5.20 ± 1.25 %, Mann-Whitney test p=0.008; Fig. 3E). The confinement area diameter was larger under latrunculin (44.87 ± 3.31 nm vs 50.72 ± 1.15 nm, t-test p= 0.008) which correspond to an increase of about 30% of the surface area. Therefore, despite a low frequency of acquisition, Pc analysis detected both the typical size of hopping compartments reported for GPI molecules and the enlargement of the confinement area size due to F-actin depolymerization.

### Using Pc to detect transient immobilizations

The number of receptors for neurotransmitters accumulated at the synapse is an important parameter setting the strength of the response and thus plays a key role in regulating neuronal function. The number of receptors depends upon the interactions that immobilize them by binding to specific scaffolding molecules. Actually, this immobilization is transient and receptors diffuse in and out of the synaptic area at unexpectedly high rates (reviewed in (27), (28)). This suggests that the receptor-scaffold interactions are short-lasting, i.e in the range of seconds.

The main ionotropic receptors at excitatory synapses are AMPA-and NMDA-type glutamatergic receptors. The number of AMPA receptors (AMPARs) fluctuates rapidly with receptors swapping between extra-and intra-synaptic areas and this dynamic accounts for the construction and plasticity of excitatory synapses (reviewed in (27),(28)). AMPARs have a preponderant role in the expression of synaptic plasticity, therefore an important question is that of their stabilisation by synaptic activity. Actually, global modifications of network activity tune the mobility of AMPARs (reviewed in (27),(28)).

Here, we compared the diffusion of AMPARs (GluA1 subunit with QDs-bound antibodies; QD-GluA1) in control conditions and after inducing synaptic plasticity. QD-GluA1 were detected with a localization accuracy of 20-30 nm and tracked at 20 Hz. The trajectories were then chopped and sorted into extrasynaptic and synaptic ones as described (29).

The stabilization periods (“immobility”) were identified by a highly confined diffusion in a small area whose size was of the order of the localization accuracy. In order to detect these events and given the localization accuracy, we set a threshold of *Pc_thresh_* = 3300 μm^−2^ (corresponding to a confinement area size with a diameter of ~35 nm) and time threshold of 0.375 s (5 time points at 20 Hz). During the recording session, *Pc* values varied displaying no, one or more stabilization events in synapses (exemplified in Fig. 4A). 35.83 ± 2.44 % of synaptic QD-GluA1 had at least one stabilization event (Fig.4B).

**Fig. 4:**
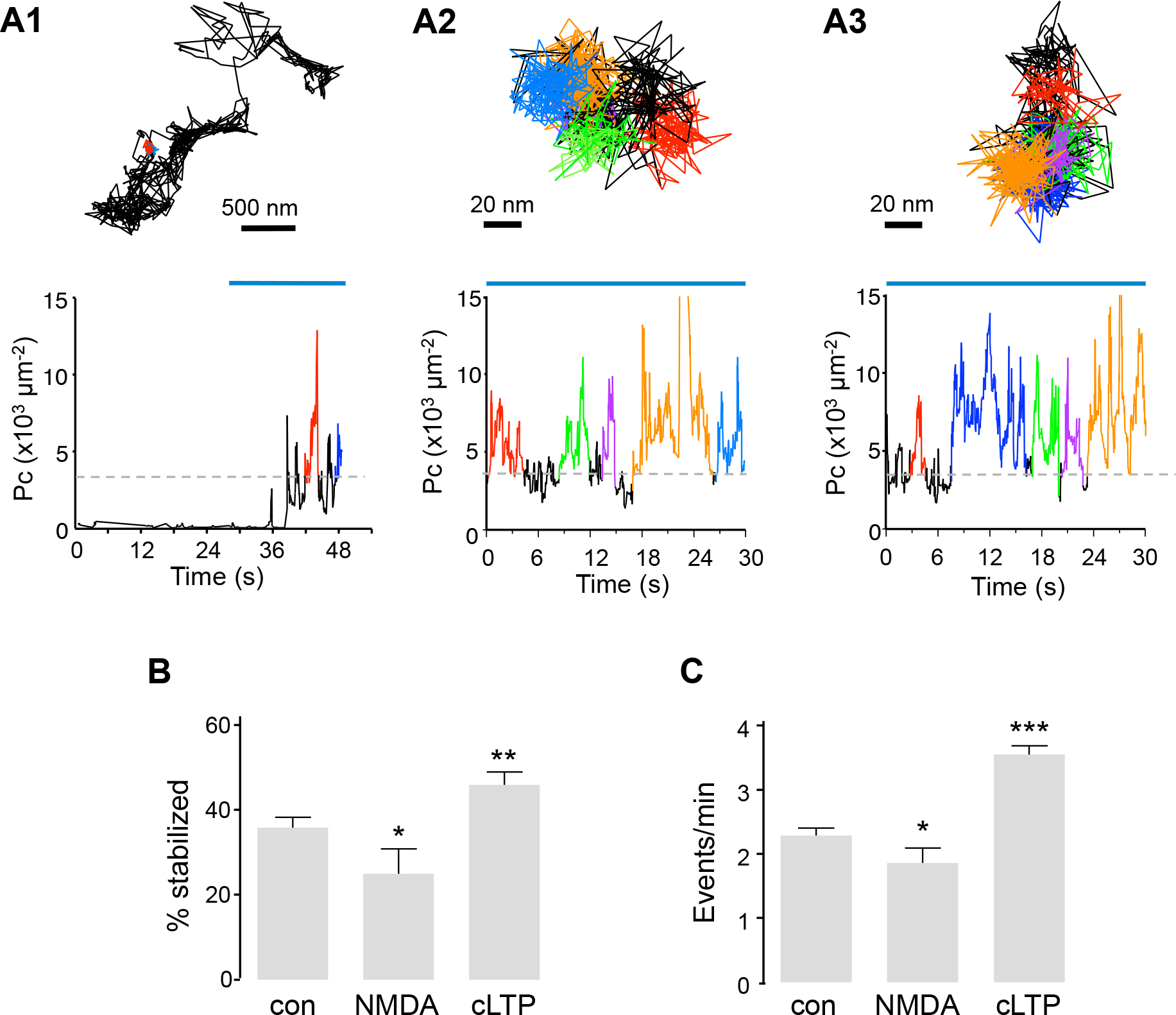
Stabilization of AMPAR analysed by *Pc* analysis. (A) Examples of *Pc* values upon time for QD-GluA1. The detected periods of stabilisation are shown in color. Top: Examples of trajectories in a dendritic spine (A1) or in synapses (A2-3). Bottom: the corresponding plots of *Pc* vs time for each trajectory. The horizontal blue line shows the synaptic localization in time. The horizontal discontinuous line shows *P_thresh_*. (B) Percentage of stabilized trajectories of QD-GluA1 in the indicated conditions (mean+/-s.e.m., n=16-48 recordings, t-test *: p<0.05, **: p<0.01). (C) Frequency of stabilization events (number of events per minute) (mean+/-s.e.m., n=331-1440 trajectories, t-test *: p<0.05, *** p<0.001).

Assuming that the stabilisation results from the first order reaction

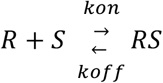

where R are the receptors and S the scaffolding molecules, we can define an effective forward binding rate *k_on_* that corresponds to the frequency of the binding events and an effective backward binding rate *k_off_* that is the reciprocal of the mean duration of the stabilization periods ((6) and references therein).

Synaptic QD-GluA1 underwent 2.29 ± 0.11 stabilization events per minute (Fig. 4C), thus indicating an effective *k_on_* = 3.81 ± 0.18 x 10^−2^ s^−1^. The duration of the stabilization events was variable (spanning between 0.375 and 50 s; the limit of our recording session), with a mean of 8.49 ± 0.32 s that corresponds to a *k_off_* of 0.11 s^−1^. In some cases events lasted the whole recording session (50 s) leading to a bias in the measurements of *k_off_*; however these long stalling periods were scarce. Despite this limitation, our effective *k_off_* was similar to the one obtained by Czöndor et al. (30) by combining FRAP and SPT data together with simulations (0.1 s^−1^).

On the other hand, the effective *k_on_* obtained here was lower than the value obtained by Czöndor et al. (30) (1.5 s^−1^), who calculated *k_on_* by fitting the reduction of global diffusion of AMPARs along the maturation of synapses. However, the *k_on_* calculated in this way reflects the random walk of receptors outside synapses and their probability to find synapses. Our effective *k_on_* was calculated only for receptors that are already in synapses and thus corresponds to the probability to find and to bind to a scaffolding site.

The low value of *k_on_* suggests that these sites were not readily accessible and/or that the affinity of receptors for their scaffolding molecules was low. In agreement with this, synaptic QD-GluA1 were immobilized only during 11.50 ± 0.66 % of the whole time that they spent in synapses. However, AMPARs often displayed more than one stabilization event (Fig. 4A2-3), therefore they were likely to undergo multiple short-lasting binding-unbinding events. The molecular crowding at the postsynaptic membrane may contribute to the successive trapping events by reducing the escape of receptors off synapses (7, 31, 32). Our results suggest that the proportion of receptors considered as “immobile” by *MSD* and *D* analysis, or the “stable fraction” obtained by FRAP (~50%,(33, 34)) are receptors that do not exit the synapse during the recording time although they are not necessarily immobilized during the whole recording period. *Pc* analysis revealed that immobilization events could be multiple and short-lasting, which could help synapse to rapidly exchange receptors with the extrasynaptic area, i.e. to replace desensitized receptors by naïve ones (34, 35).

Synaptic plasticity mechanisms rely on changes in the number of AMPARs, which is increased or decreased during long term potentiation (LTP) and long term depression (LTD), respectively (reviewed in (36)). The accumulation of synaptic AMPAR is decreased by NMDA application (referred to as a model for chemical LTD, (9)) whereas chemical LTP (cLTP) protocols induce the enrichment of GluA1-containing AMPARs (10).

Neuronal cultures were challenged with a cLTP protocol or NMDA application in order to modify the stabilization of QD-GluA1 at synapses. As expected, both treatments had opposite effects on the stabilization of receptors, affecting both the percentage of stabilized QD-GluA1 (control = 35.83 ± 2.44 %, NMDA= 24.94 ± 5.88 % Mann-Whitney test p=0.041, cLTP= 45.95 ± 2.97 % Mann-Whitney test p=0.007, Fig. 4B) as well as the number of immobilization sequences per minute (control = 2.29 ± 0.11 events/min, NMDA= 1.85 ± 0.23, Mann-Whitney test p=0.025, cLTP= 3.55 ± 0.13, Mann-Whitney test p<0.0001, Fig. 4C). Thus, in agreement with the reported reduction of AMPAR amount in synapses (refs in (10)), NMDA application decreased their trapping probability. Conversely, cLTP protocol increased their trapping probability, what is consistent with the increased number of receptors in excitatory synapses induced by this treatment (refs in (9)).

Overall, the *Pc* approach presented here allowed the identification of transiently stabilized trajectories, thus providing a better and new characterization for the sub-synaptic diffusion of receptors and the computation of effective kinetic parameters of scaffolding interactions. Furthermore, the *Pc* analysis provided the spatial localization of the stabilization events. This approach could then be combined with super-resolution microscopy of post-synaptic scaffold molecules in order to investigate fluctuations of the trapping of receptors (37) with respect to the nano-organisation of the postsynaptic scaffold (38).

### Application of Pc analysis to short trajectories

Until recently, single molecule studies were restricted to only a few spatially isolated molecules sparsely labeled on living cells. A number of approaches have emerged that generate reconstructed images of single-molecule localizations at high density, such as sptPALM or universal point-accumulation-for-imaging-in-nanoscale-topography (uPAINT, (12)). The drawback of these techniques is that the resulting trajectories are short due to the rapid photobleaching of fluorophores. Indeed, short trajectories constrain the time window used to calculate *Pc*, thus lowering its statistical power. In addition, short trajectories do not display a confined behaviour if they are not long enough to sense the limits of the confined area. Moreover, the relatively poor signal-to-noise ratio of available fluorophores introduces extra noise reducing the localization accuracy. Therefore, we questioned if *Pc* analysis could be used in case of short trajectories. Thus, we simulated trajectories with lengths of 5 or 10 time points, being confined or not in areas of different sizes. We took into account localization accuracies related to common fluorophores (Fig. 5A1-2). *Pc* revealed confinement at different levels in the longer trajectories (Fig. 5A1). As expected, the localization accuracy had an important effect on *Pc* values. As a rule of thumb, the minimum size of confinement that could be detected was twice the localization accuracy. In the case of shorter trajectories, *Pc* values were highly scattered (Fig 5A2). Thus, if the localization accuracy is good enough (30 nm or less), the *Pc* analysis on trajectories of less than 10 points could be used only to detect immobility (L<60 nm in this case).

**Fig. 5:**
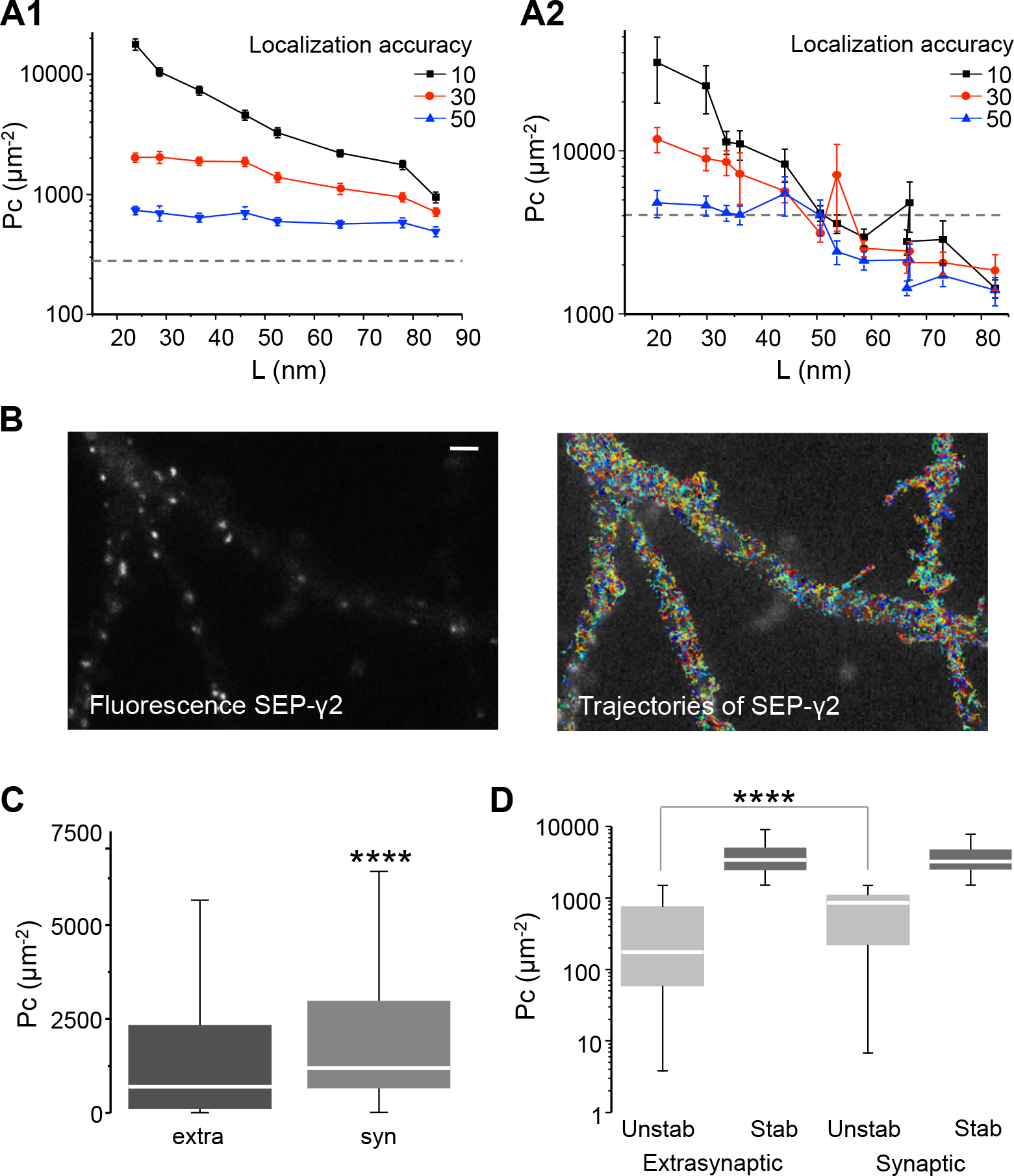
*Pc* analysis applied to short trajectories. (A) Mean *Pc* values 10 time points-long (A1) or 5 time points-long (A2) simulated trajectories confined in areas of the indicated sizes (L) (mean+/- s.e.m., n=100 trajectories in each case). Simulations took into account different localization accuracies. The horizontal broken line: P95 value of *Pc* distribution of random walk trajectories. (B) Fluorescent image of SEP-γ2 displaying synaptic puncta (right) overlaid with trajectories (each trajectory in a different color) of nanobodies-labelled SEP-γ2 (left). Bar: 1 μm. (C) Distribution of *Pc* (median, box: 25%-75% IQR, whiskers: 5%-95%) for SEP-γ2 trajectories in (syn) and out (extra) synapses. Mann-Whitney test ****: p<0.0001, n synaptic = 427, n extrasynaptic = 4319. (D) Distribution of *Pc* (median, box: 25%-75% IQR, whiskers: 5%-95%) values of trajectories in C, after sorting into stabilized (Stab, *Pc*>1241 μm ^−2^) and unstabilized (Unstab) (Mann-Whitney test ****: p<0.0001, n= 199-2838).

To test the suitability of Pc for experimental data, we performed uPAINT acquisitions to analyze the diffusion of GABA receptors on the surface of hippocampal neurons (Fig. 5B). uPAINT consists in recording single-molecule trajectories that appear sequentially on the cell surface upon the continuously labeling of the molecules of interest (12). Membrane molecules are labeled with fluorescent ligands that are diluted in the cell medium to the adequate concentration, in order to obtain a sparse labeling at each time point. Only ligands bound to membrane molecules are considered, by excluding fluorescent molecules freely diffusing in solution (see materials and methods). uPAINT provides massive amounts of trajectories that can be used to create “maps” of the diffusive state of membrane molecules. Neurons were transfected with SEP-tagged γ2 subunit of GABA_A_ receptors (SEP-γ2, (8),(29)) which could be tracked using extracellular labelling with ATTO647N-coupled nanobodies against GFP. Actually, GABA_A_ receptors are pentamers of at least three different subunit isoforms, being immobilized or not at synapses depending on their subunit composition ((39) and references therein). The majority of γ2 subunit-containing receptors are accumulated in synapses, however γ2-GABA_A_R that include also the α5 subunit are not stabilized by the synaptic scaffold ((29) and references therein). In fact, α5-containing GABA_A_R are trapped outside synapses by interactions with radixin (40), thus some SEP-γ2 labels receptors can be stabilized either in or out synapses. Fluorescent images (10000-20000 consecutive frames) were recorded at 33 Hz with a localization accuracy of ~30 nm. The obtained trajectories had a median length of 12 time points (first and third quartile 8 and 19, respectively). We kept only trajectories with at least 10 time points, calculating *Pc* on a sliding window of 10 points. Only one *Pc* value was extracted for each trajectory: for trajectories longer than 10 points, *Pc* was averaged among all the calculated values.

Synapses were identified by the fluorescent spots of SEP-γ2 (Fig. 5B), and trajectories were sorted into extrasynaptic and synaptic as previously (29). As expected from the enrichment of GABA_A_R in synapses, the *Pc* values distributed around higher values for synaptic SEP-γ2 than for extrasynaptic ones (Fig. 5C). In order to sort “stabilized” and “unstabilized” trajectories, given our localization accuracy, we set *Pc_thresh_* to 1241 μm^−2^ (corresponding to a confinement area size with a diameter of ~60 nm) and *t_thresh_* to 0.33 s (10 time points at 30 Hz). With these settings, the proportion of stabilized trajectories was higher in synapses (46.60%) than at extrasynaptic locations (34.29%). In agreement with the increased confinement of diffusion in synapses reported previously (29), synaptic unstabilized trajectories displayed higher confinement than unstabilized extrasynaptic ones (Fig. 5D). In conclusion, even if trajectories obtained with uPAINT were indeed too short to detect transitions between different diffusive states, *Pc* analysis successfully detected different levels of confinement and stabilization of SEP-γ2 GABA_A_R in and out synapses.

## Conclusions

The *Pc* parameter defined here allows the detection and quantification of transient confinement sequences. The main advantages of this analysis approach are: 1) simple implementation and rapid calculation, 2) calculation from a sliding window thus allowing the detection of transient changes and their localization, and 3) independency from *MSD* & *D* calculation, thus it can be used to compare molecules or situations with different global diffusivity (i.e. comparison of confinement between molecules with different mobility, or analysis of diffusion-trapping in crowded environments such as synapses).

Above all, the *Pc* analysis allows the detection of stabilization events along trajectories thanks to their sensitivity to confinement within small (<50nm) areas. Thanks to this, it is then possible to derivate the effective kinetic constants of the molecular interactions implicated in the stabilization even if interactions are weak and transient. This type of short-lived interactions is difficult to grasp with classical bulk methods (such as co-immunoprecipitation, mass spectrometry or isotitration calorimetry, (37)) although it can be an important parameter being regulated during cellular processes such as synaptic plasticity.

## Acknowledgements

We would like to thank Professor Stuart Edelstein for his constant support and critical review of this manuscript. We thank the IFM imaging platform for technical support. This work was supported by the Agence Nationale de la Recherche “Synaptune” (Programme blanc, ANR-12-BSV4-0019-01), the ERC advanced research grant “PlasltInhib”, the program “Investissements d’Avenir” (ANR-10-LABX-54 MEMOLIFE and ANR-11-IDEX-0001-02 PSL* Research University), and the Institut National de la Santé et de la Recherche Médicale (INSERM). L.H. was supported by the Ministère de la Recherche et de l’Enseignement Supérieur. LW was supported by the China Scholarship Council (CSC, fellowship No. 2011614094). Authors declare no conflict of interest.

## Author Contributions

MR designed research, performed research, contributed analytic tools, analyzed data and wrote the manuscript. LW performed research. LH performed research and analyzed data. SL contributed with materials and equipment. AT designed research and wrote the manuscript.

---

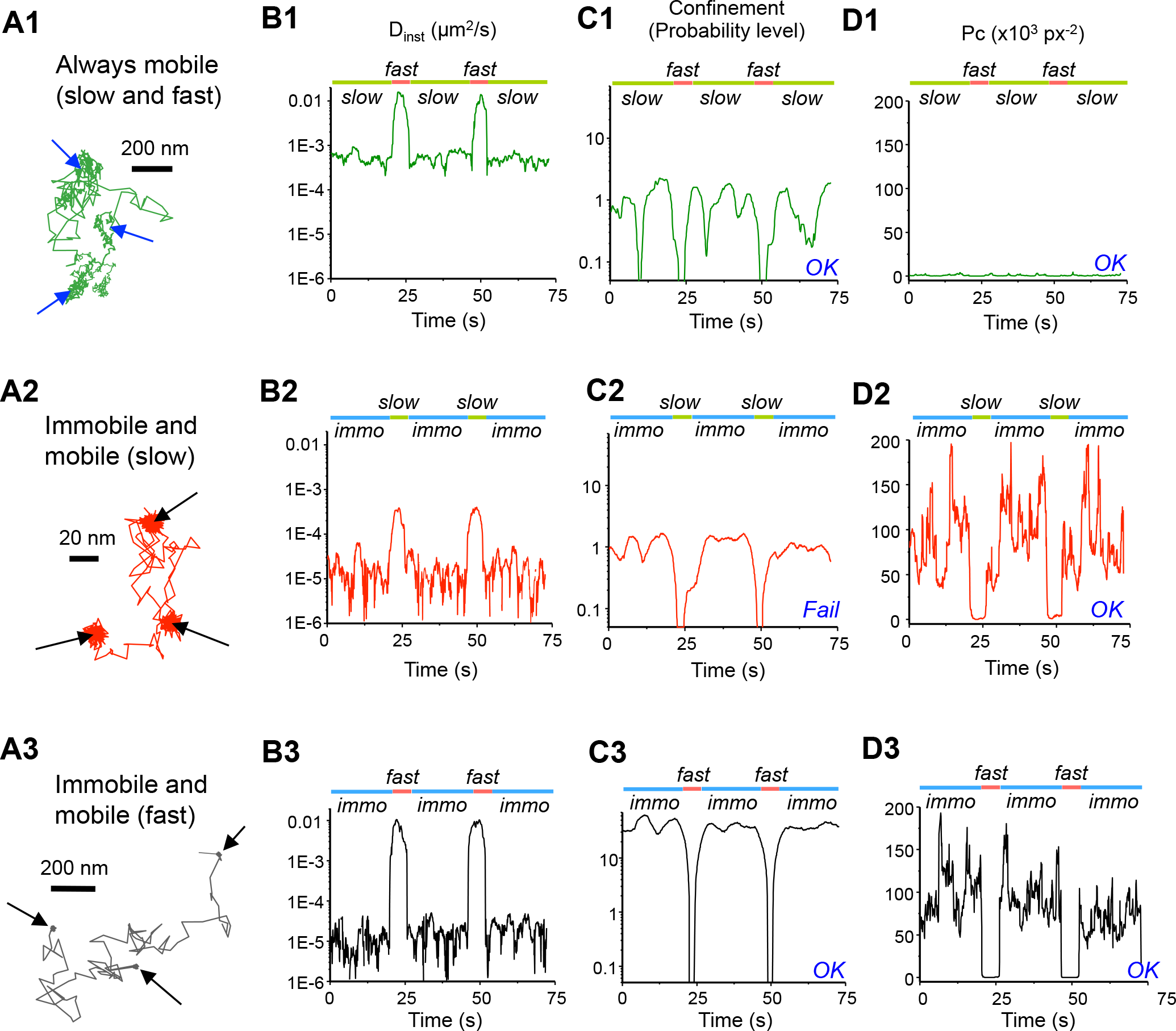

**Fig. S2:**
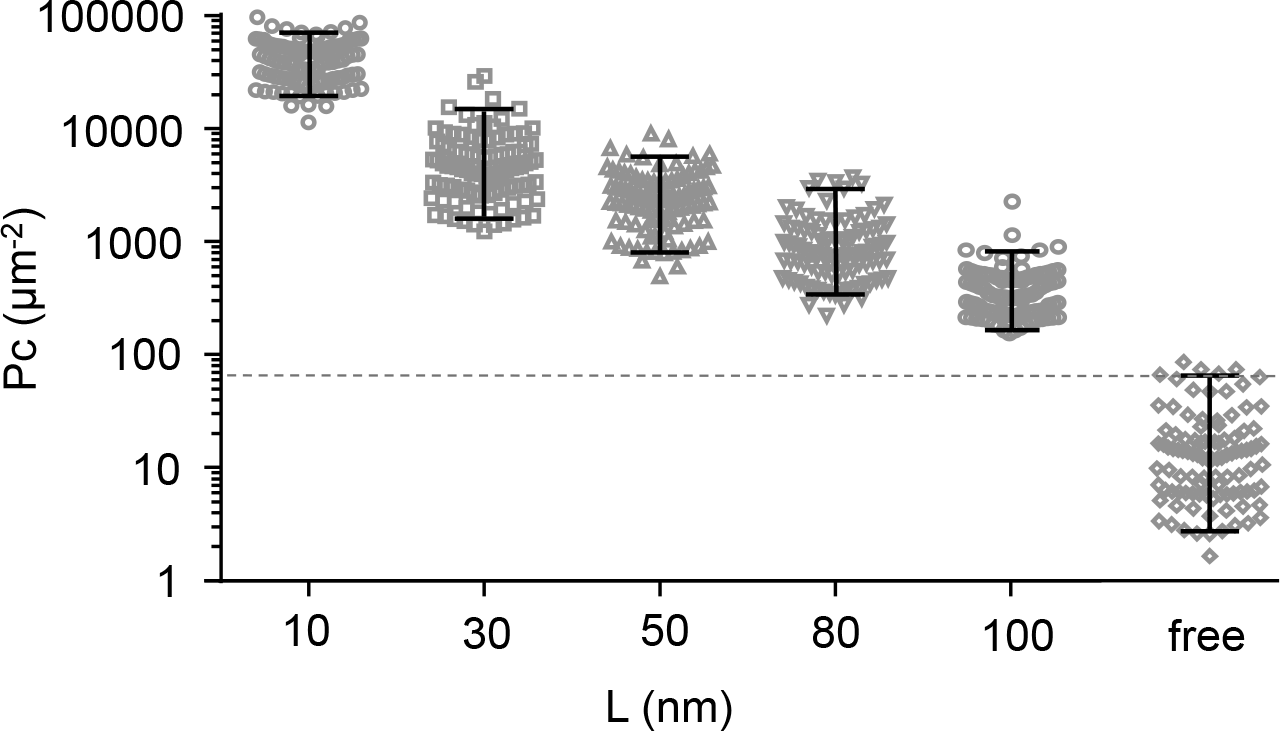
Distribution of *Pc* values on 30 time points-long simulated trajectories confined in areas of the indicated sizes (L) or not confined (free) (bars: P5 and P95 of the distribution, n=100 trajectories in each case). Simulations were done taking into account a localisation accuracy of 10 nm. The horizontal broken line corresponds to the P95 value of *Pc* distribution of random walk trajectories (67 μm^−2^).

